# Culture-Independent Detection and Identification of *Leptospira* Serovars

**DOI:** 10.1101/2022.06.24.497575

**Authors:** Michael A. Matthias, Aristea A. Lubar, Shalka S. Lanka Acharige, Kira L. Chaiboonma, Nicholas N. Pilau, Alan S. Marroquin, Dinesha Jayasundara, Suneth Agampodi, Joseph M. Vinetz

## Abstract

Pathogenic *Leptospira*, the causative agents of leptospirosis, comprise >200 serotypes (called serovars). Most have a restricted reservoir-host range, and some, e.g., serovar Copenhageni, are cosmopolitan and of public health importance owing to their propensity to produce severe, fatal disease in humans. Available serotyping approaches—such as multi-locus sequence typing, core genome sequence typing, pulsed-field gel electrophoresis, and the cross-agglutination absorption test—are tedious and expensive, and require isolation of the organisms in culture media—a protracted and incredibly inefficient process— precluding their use in prospective studies or outbreak investigations. The unavailability of culture-independent assays capable of distinguishing *Leptospira* serotypes remains a crucial gap in the field. Here, we have developed a simple yet specific real-time qPCR assay—targeting a *Leptospira*-unique gene encoding a putative polysaccharide flippase—that provides intra-species, serotype-defining (i.e., epidemiologically useful) information, and improves upon the sensitivity of preferred *lipL32*-based qPCR-based diagnostic tests. The assay, dubbed RAgI (“rage one”), is <u>r</u>apid and <u>a</u>ffordable, and reliably and specifically detects <u>g</u>roup <u>I</u> pathogenic *Leptospira* in culture, serum and urine, with no detectable off-target amplification—even of the genetically related but low virulence group II pathogenic (formerly “intermediate”) or non-pathogenic *Leptospira*. It retained 100% diagnostic specificity when tested against difficult sample types, including field-collected dog urine-samples and environmental samples containing varied and complex microbial species-consortia. And holds considerable promise in the clinical setting, and for routine epidemiological and environmental surveillance studies.

## Introduction

Leptospirosis is a sometimes-lethal, waterborne infectious disease caused by pathogenic *Leptospira* (1–4). Humans are usually infected via contact with contaminated surface water or damp soil, but occupational exposure in kennels, slaughterhouses and animal farms, does occur—albeit relatively infrequently. Rodents are notorious ‘reservoir’ hosts, but virtually any terrestrial species, including pets and domesticated animals, can carry and therefore disseminate *Leptospira*. Transmission is presumably more common in warm, wet locales, but disease is cosmopolitan with a global burden comparable to cholera and typhoid fever (5, 6). Epidemics grow more frequent, made worse by changing weather patterns and increased overcrowding in urban slum communities (5–7). Even so, the scarcity of epidemiological data— well-curated or otherwise—insufficient public health awareness, extensive evidence gaps and a lack of adequate diagnostic tests that, collectively, artificially depress incidence rates, indicate that the public health impact of leptospirosis is greatly underappreciated. Yet, despite several well documented limitations—mainly their inability to provide definitive results soon enough for clinical interventions to be effective (8)—<u>A</u>nti<u>b</u>ody-based diagnostic <u>T</u>ests (ABTs), e.g., the <u>M</u>icroscopic <u>A</u>gglutination <u>T</u>est (MAT) or IgM-based ELISAs remain the norm, only worsening the public health outlook.

Newly developed leptospirosis diagnostic tests have tended to focus instead on antigen detection. Most are <u>n</u>ucleic <u>a</u>cid <u>a</u>mplification-based diagnostic tests (NAATs) that target various leptospiral genes and are designed to provide an early diagnosis of leptospirosis (9–14). In practice, many are plagued by context-related problems—particularly a low diagnostic sensitivity when epidemiological information is lacking—so few are generalizable. Leptospirosis NAATs-du-jour target *lipL32* (15–17), a taxonomically restricted gene found almost exclusively in *Leptospira. Leptospira* alleles are distinctive and are found only in pathogenic species, ergo *lipL32*-based assays are specific for medically important *Leptospira*. They tend to be remarkably consistent (dynamic range of 10^7^ – 10^0^ (15)) but produce somewhat unpredictable results with non-*L. interrogans* leptospiral species, owing to the high sequence diversity of the targeted gene. Despite these recognized benefits, *lipL32*-based assays are not yet routine in clinical or public health settings, presumably because the best options employ fluorescent probes inflating setup and operational costs, which renders them ill-suited to the low-tech’ realities of most leptospirosis endemic regions. To expand access, cheaper PCR-based assays have been developed (18) but these by their nature suffer from inferior (i.e., higher) detection limits and are tedious compared to qPCR-based alternatives. Regrettably, *lipl32*-based NAATs provide little insight WRT infecting *Leptospira* species or serotype. Accordingly, they are best suited to the clinical setting where an early diagnosis is paramount.

Some leptospirosis NAATs allow species identification (ID), e.g., *OmpL1*-PCR (19), but a universal limitation of these assays has been their inability to provide serotype information. Serovar ID remains both clinically and epidemiologically useful, as some serovars, e.g., Copenhageni (shorthand notation, *sv*Copenhageni), are especially virulent, and many have preferred reservoirs, e.g., *sv*Copenhageni for rats, and *sv*Canicola and *sv*Hardjo for dogs and cattle, respectively (1). Reputed ‘Gold Standard’ approaches for serotyping *Leptospira* strains, such as the Cross Adsorption Agglutination Test (CAAT), and its modern equivalent utilizing panels of cross-reacting <u>m</u>onoclonal <u>a</u>nti<u>b</u>odies (mAbs), are complicated, expensive and impractical. <u>N</u>ucleic <u>A</u>cid <u>B</u>ased <u>S</u>erotyping (NABS) methods, such as <u>M</u>ulti-<u>L</u>ocus <u>S</u>equence <u>T</u>yping (MLST) (20, 21), <u>M</u>ulti-<u>L</u>ocus <u>V</u>ariable <u>N</u>umber <u>T</u>andem <u>R</u>epeat Analysis (MLVA) (22) and, more recently, <u>c</u>ore <u>g</u>enome <u>M</u>ulti-<u>L</u>ocus <u>S</u>equence <u>T</u>yping (cgMLST) (23) are useless clinically due to their reliance on culture isolation (as do all current *Leptospira* serotyping methods), and impractical in most epidemiological contexts. Indeed, several NABS-rooted schemes have been proposed and abandoned for varied reasons but primarily their inconsistent performance with non-*L. interrogans* species. Consequently, development of inexpensive, easily deployable, culture-independent, antigen-detection tests capable of distinguishing *Leptospira* serovars is paramount.

Here, we report on the development and validation of the first <u>r</u>apid <u>a</u>ffordable <u>g</u>roup <u>I</u>-specific (RA*g*I) NAAT capable of reliably distinguishing *Leptospira* serotypes in diverse sample types, obviating requirements for culture isolation. The assay, which we’ve dubbed the leptospirosis 11108 **RA*g*I** (or “rage-one”) test, targets LIC_RS05715 (old locus_tag: LIC_11108)—a distinctive, single-copy, leptospiral core gene encoding a putative oligosaccharide flippase (NCBI protein ID: <u>AAS69715.1</u>; UniProtKB ID: <u>Q72TB3</u>)—and allows joint inference of *Leptospira* species and serovar. The data presented here provide crucial proof-of-concept validation for and should spur development of conceptually related leptospirosis <u>s</u>erotyping NAATs, (i.e., sNAATs).

## Materials and Methods

### Primer design

For real-time qPCR-based serovar identification (ID), genes comprising the core *Leptospira* genome were first defined using the Pan-Genome Ortholog Clustering Tool, PanOCT (24), and those predicted to encode proteins participating in LPS assembly or export, e.g., O-antigen polymerases (*rfbY*), O-antigen/polysaccharide flippases, *rfbX* and LIC_11108, and the *Leptospira* lipopolysaccharide export system (Lpt), including *msbA, lptA* and *lptD*, were used to construct exhaustive amino-acid-guided <u>m</u>ultiple <u>s</u>equence <u>a</u>lignments (MSAs) of coding sequences (CDS) retrieved from Genbank—anticipating that these would be Serogroup- and (potentially) serovar-dependent, due to their stereospecificity (25). LIC_11108 (abbreviated 11108 hereafter) was chosen for initial evaluation based on several criteria (the most notable summarized in the preceding paragraph).

From these MSAs, hyper-variable regions were identified and targeted for amplification by primers complementary to conserved flanking sequence. As none were 100% conserved, degenerate primers 11108*f*: 5’ – TT**<u>R</u>** AA**<u>N</u>** GA**<u>R</u>** AGT ATA AAA CTT CC – 3’, and 11108*r*: 5’ – AAC GG**<u>R</u>** CT**<u>Y</u>** TTT CAA TA**<u>Y</u>** TA**<u>Y</u>** GC – 3’ (Table 1)—specific for group I pathogenic *Leptospira*—were synthesized and validated against 26 well-characterized *Leptospira* serovars. Which included common human-infecting strains isolated in Iquitos, Peru and various villages in Sri Lanka (Table 2), and those constituting the MAT reference panel recommended by the Centers for Disease Control and Prevention (CDC) (Table 2). All strains were maintained in liquid <u>E</u>llinghausen-<u>M</u>cCullough-<u>J</u>ohnson-<u>H</u>arris (EMJH) as described previously (26). Genomic DNA (*g*DNA) for qPCR (and sequencing) was prepared using the manufacture-recommended extraction protocol for Gram-negative bacteria (GeneJET™ DNA Purification Kit, Thermo Scientific™), and then amplified on a CFX96 C1000 thermal cycler (Bio-Rad). Thermal cycling was followed by post-amplification high-resolution melt-curve analysis using pre-installed Precision Melt Software (purchased separately). Conscious of our intended longer-term development goals, we opted for a SYBR™ Green-based NAAT—which can be adapted for TaqMan-based assays—and evaluated the performance of the empirically optimized assay using more affordable hardware/software options: CFX96 **w/**default melt-curve analysis (i.e., without Precision Melt Software) and on affordable hardware—specifically a modestly priced four-channel <u>m</u>agnetic induction thermal <u>c</u>ycler (MIC) manufactured (and sold) by Bio Molecular Systems. The MIC is Wi-Fi-enabled (ergo networkable), portable (dimensions, 15 cm x 15 cm x 13 cm / weight, 2.1 kg), rapid (completing typical 40 cycle runs in under 30 min), and remarkably precise (<0.1°C resolution).

**Table 1:**
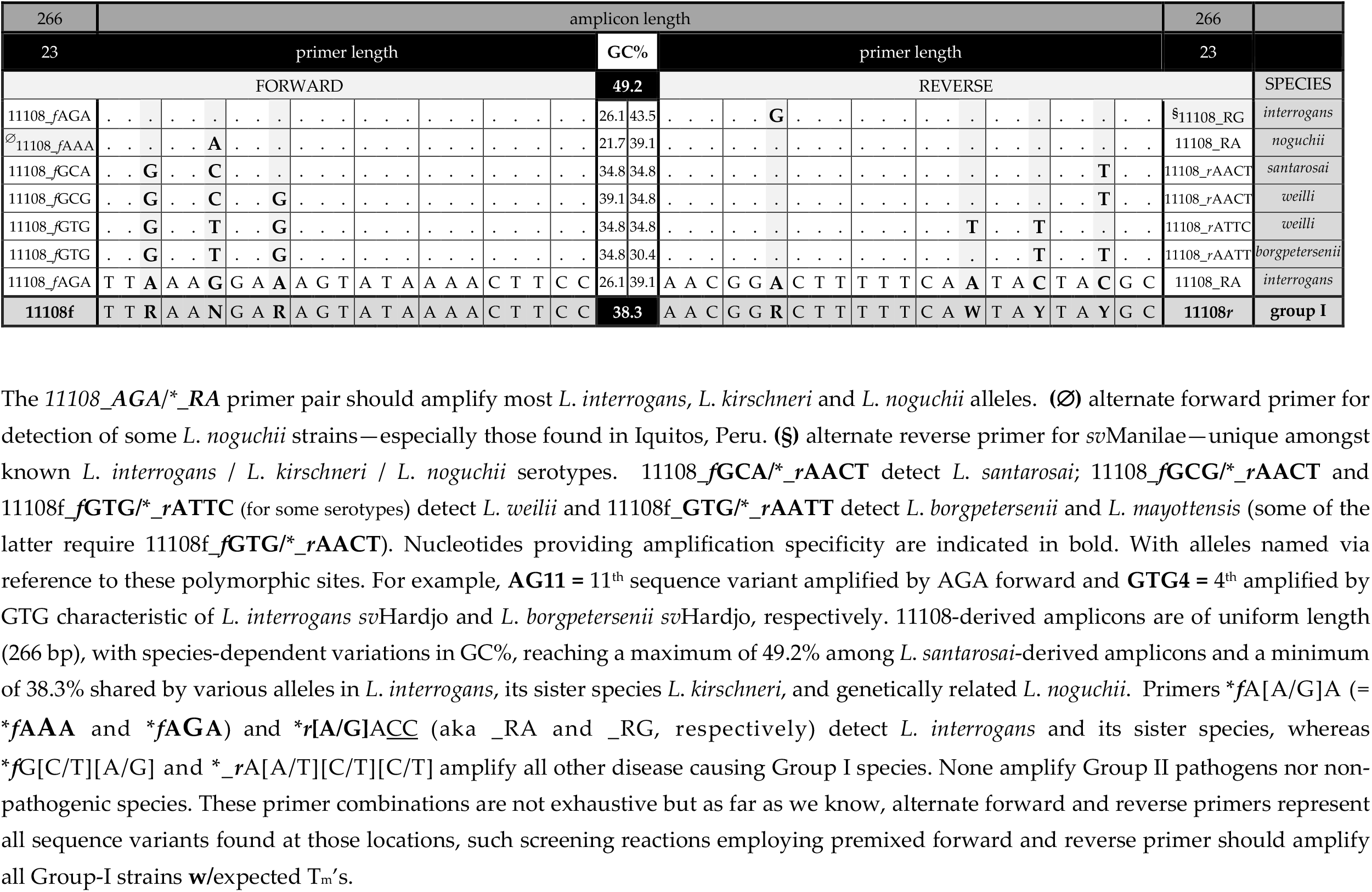
11108 qPCR assay, forward and reverse primer pairs

**Table 2:**
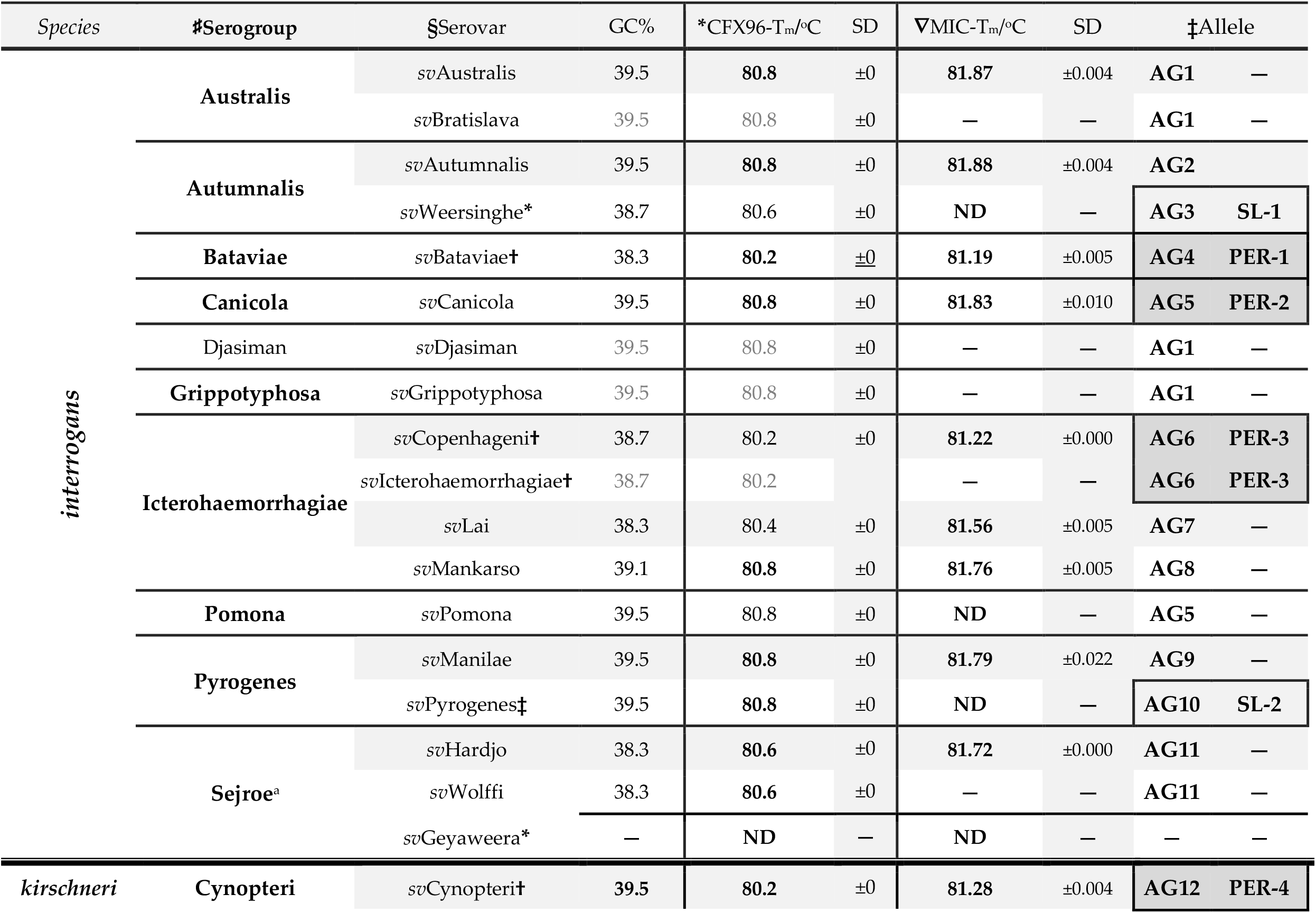
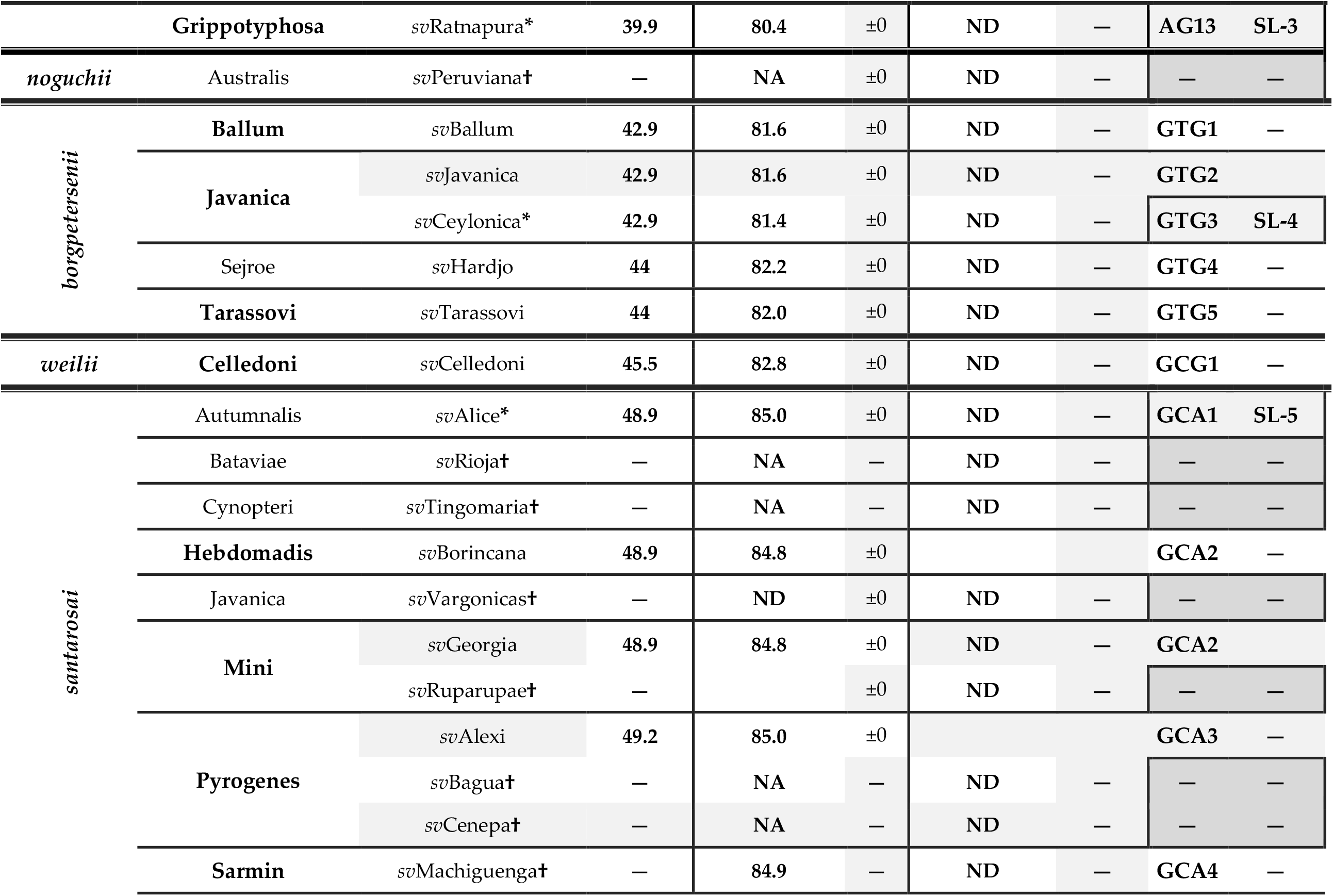
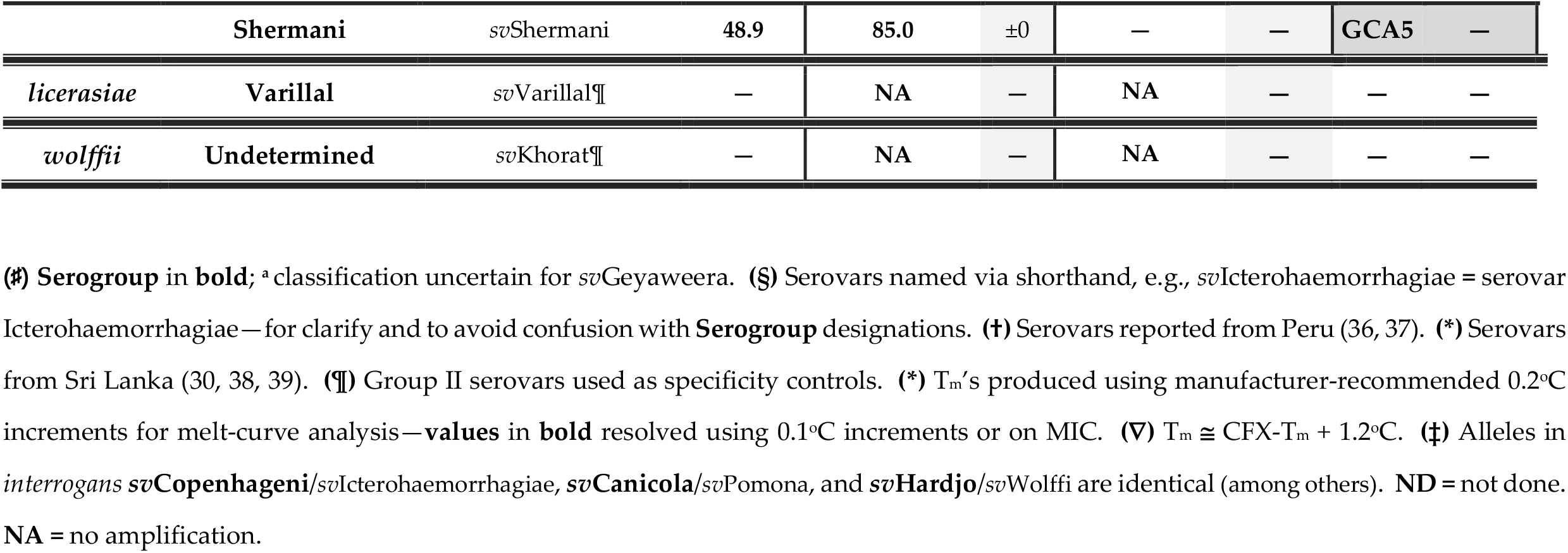
Serotype-Characteristic T_m_’s produced by various *Leptospira* strains, including isolates from Peru and Sri Lanka

### Validation and optimization of *11108*-based assay

Pilot experiments were done using degenerate primers to establish baseline assay conditions before re-amplification and optimization with de-convoluted species-appropriate primers (Table 1) to produce characteristic serovar-dependent melt temperatures that we refer to as diagnostic T_m_ for simplicity. Reaction mixes consisted of the following: 10 *µ*L of Precision Melt Supermix, 2 *µ*L of template *g*DNA, and 4 *µ*L of both forward and reverse primers, yielding final (empirically optimized) primer concentrations of 0.4 *µ*M per 20 *µ*L reaction. The finalized thermal cycling profile consisted of a 3 min hold at 95°C (denaturation), followed by 44 cycles of 95°C for 10s, 62°C for 30s (empirically optimized annealing temperature) and 72°C for 1 min (extension), and a final hold at 72°C for 7 min to ensure full extension of PCR product. Amplification was followed by melt-curve analysis with fluorescence measured at 0.2°C increments. All samples were run in triplicate, and runs were repeated at least thrice to produce a minimum of nine data points/per sample. Cq values of ≤ 40 cycles were considered positive, but post-cutoff amplification (Cq > 40) with appropriate melt peaks were recorded, and later verified by sequencing. Milli Q water was run in parallel, in triplicate, as a no template control (NTC). *g*DNA from *L. licerasiae* **Varillal** *sv*Varillal (archetype of group II, comprising low virulence *Leptospira* species) and the genetically related *L. wolffii sv*Khorat (Table 2) and pre-mixed *g*DNA from Gram-negative and Gram-positive bacteria, *Citrobacter rodentium* (ATCC DBS 100), *Escherichia coli* (EPEC E2348/69), *Clostridium difficile* (ATCC BAA-9689) and *Staphylococcus aureus* (MARSA, USA 300), were used as negative controls (to assess non-specific “off-target” amplification). For each serovar, a mean T_m_ was calculated after first identifying and then removing outlier data points via robust <u>r</u>egression and <u>out</u>lier removal (ROUT)(27) analysis (Q=10%, Prism v7.0a). Serovar-characteristic T_m_’s were defined via comparison of sample-means by nonparametric *t-*test (Prism v7.0a).

qPCR results were verified by conventional Sanger sequencing of cloned amplification products (TOPO™ TA Cloning™ Kit, Invitrogen). Twenty-five randomly chosen colonies were sub-cultured into LB broth, plasmids were isolated using a QIAprep® Spin Miniprep Kit (Qiagen) then sent for sequencing. Consensus sequences were aligned using the MAFFT <u>m</u>ultiple <u>s</u>equence <u>a</u>lignment (MSA) algorithm with default parameter settings. Individual consensus sequences were also BLAST*ed* to confirm specificity (28). For serovars consistently yielding inexact or a quasi-stable T_m_’s, we sequenced 10 randomly selected clones producing six reads per clone (for a total of 60), and then compared clone-specific consensus sequences to identify variants.

### Detection limit and cross-species amplification of de-convoluted (i.e., species-defined) 11108 primer sets

To determine the detection limit of individual species-defined primer sets, amplicons from *interrogans sv*Lai, *santarosai sv*Shermani and *borgpetersenii sv*Tarassovi were TA cloned to construct reference(-gene-containing) plasmids. After routine Minipreps, purified (reference) plasmids were quantified (individually) by fluorometry (Qubit™, Thermo Fisher Scientific), and copy number estimated. Reference plasmids were serially diluted in either PBS buffer or heat-inactivated <u>n</u>ormal <u>h</u>uman <u>s</u>erum (NHS) and re-extracted using the QIAamp® DNA Blood Mini Kit (Qiagen) per recommended protocol for serum. The detection limit was defined as the minimum number of real-time-qPCR-detectable copies producing the expected T_m_ among all technical replicates. Since 11108 genes occur once per *Leptospira* genome, copy numbers could be expressed as <u>g</u>enome <u>e</u>quivalents (GEs). Reference plasmids were used as template in amplifications examining the propensity of de-convoluted primer sets for cross-species amplification.

### Serovar resolution via high-precision magnetic induction thermal cycling

Serovars that could not be distinguished (based on T_m_) by block-based thermal cycling (CFX96) were re-analyzed via high-precision MIC. Reaction mixes and thermal cycling profile were as described above, but for post-amplification melt-curve analysis absorbance readings were taken at increments of 0.025°C exploiting the superior precision of MIC. Negative controls and NTCs were run in triplicate, in parallel, as described above. And as before, outliers were removed by ROUT analysis before comparison of serovar-associated T_m_ by nonparametric *t*-test. Data were visualized via a box-and-whisker plot.

### Reliability, accuracy, sensitivity and specificity of *11108*-based assay

To evaluate the accuracy of our ***11108***-based real-time qPCR assay (and thus, its potential epidemiological utility), we analyzed 45 *Leptospira* isolates obtained over a four-year period from febrile patients, domesticated animals and rats in Iquitos and 12 (of 25) serum-isolates from hospitalized patients in Sri Lanka, assessing agreement among T_m_ ‘s, PFGE- and CAAT-derived serovar assignments (Table S1), and paired T_m_’s, respectively (Table 3). Outlier removal, statistical comparisons and data visualization were as described above. Disagreements were resolved by Sanger-sequencing amplified products. Assay sensitivity and specificity were determined using canine urine-samples having associated culture results, which were sourced from a study of canine leptospirosis in Sokoto, Nigeria (29) (as well as acute serum-samples from a prospective study of human leptospirosis in Sri Lanka (30)). Because *Leptospira* species and serovars in Sokoto and study sites in Sri Lanka are not well-characterized, urine, and serum-samples were amplified using premixed, species-appropriate primers.

**Table 3.**
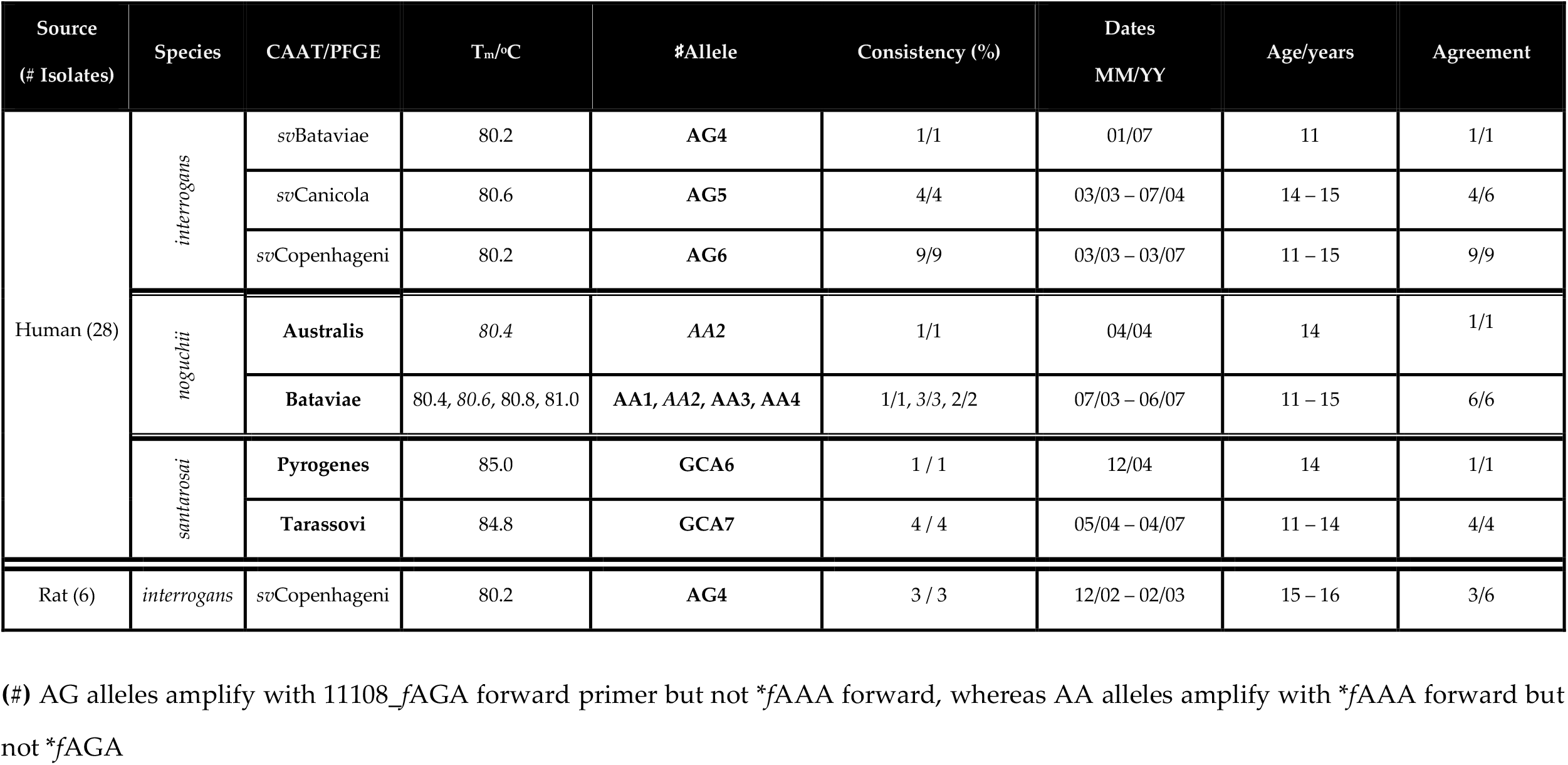
*Leptospira* strains isolated over a five-year period (2002 – 2007) from humans and peri-domiciliary rats in Iquitos, Peru. CFX96-derived T_m_ and relevant serotyping info are indicated (minimum of three replicates)

For qPCR, gDNA was purified from 84 urines comprising 40 culture positives and 44 randomly selected culture negatives. *g*DNA from *sv*Manilae was used as a positive control and as before, premixed *g*DNA from four bacterial species was used as a negative control. Samples were also assayed by qPCR amplification of *lipL32* (*lipL32_178f*: 5’ – TCT GTG ATC AAC TAT TAC GGA TAC – 3’, and *lipL32_419r*: 5’ – ATC CAA GTA TCA AAC CAA TGT GG – 3’), specific for virulent group I pathogenic *Leptospira* species. The (previously) empirically optimized *lipL32*-based thermal cycling profile consisted of the following: 3 min denaturation at 95°C, 45 amplification cycles (95°C for 10s / 62°C for 30s / 72°C for 60s), 7 min extension at 72°C followed by post amplification melt-curve analysis.

Because culture is inefficient (<10% success rate, in our hands), 11 presumed false positives (culture negatives that were positive by at least one real-time qPCR assay) were verified by <u>s</u>hotgun <u>m</u>etagenomics <u>s</u>equencing (SMS). All were positive by 11108-based real-time qPCR, and four were positive by *lipL32* qPCR (not shown). Each was sequenced in independent paired-end runs—two samples were deemed of poor quality based upon visual inspection of the raw data and were abandoned. Of the initial 84 samples, complete data were produced for 80: **47** verified positives (including real-time qPCR-only positives) and **33** negatives. Assay results were used to construct a contingency table to assess sensitivity and specificity (Table 4).

**Table 4.** Comparison of the sensitivity and specificity of 11108 and *lipL32* qPCR assays

|  | Sensitivity | Specificity |
| --- | --- | --- |
| *11108* | 91.4% | 100% |
| *lipL32* | 68.1% | 100% |

## Results

### Validation and optimization of *11108*-based assay

For most species, the optimized 11108-assay had a detection limit of 10 GEs with expected T_m_ regardless of diluent (i.e., PBS or NHS). But to achieve this level of detection with *santarosai*-specific primer pairs, extended runs (> 40 ≤ 45 cycles) were required or 10^2^ – 10^3^ otherwise. As summarized in Table 2, at 0.2°C-resolution, T_m_’s ranged from 80 – 85°C. Negative controls and NTCs were invariably negative, confirming specific detection of group I pathogenic *Leptospira*. Intra- and inter-run T_m_’s were remarkably consistent (***σ*** = 0.0°C) among replicates, with T_m_’s defined for each allele (Table 2). At the manufacturer-recommended resolution of 0.2°C, nine *interrogans*/*kirschneri* alleles (16 serovars) produced only four T_m_’s, ranging from 80.2 – 80.8°C. With a T_m_ of **80.2±0.0°C** being characteristic of *sv*Bataviae, ***sv*Copenhageni/***sv*Icterohaemorrhagiae (where ‘**/**’ indicates 100% sequence identity of associated amplicons), and *kirschneri* ***sv*Cynopteri**; **80.4±0.0**°C of ***sv*Lai**; **80.6±0.0**°C of ***sv*Hardjo-prajitno/***sv*Wolffi, *sv*Mankarso and *sv*Manilae; and **80.8±0.0°C** of ***sv*Australis/***sv*Bratislava*/sv*Djasiman/*sv*Grippotyphosa/*sv*Pyrogenes, ***sv*Autumnalis**, and ***sv*Canicola/***sv*Pomona. Nonetheless, several important serovars, specifically ***sv*Copenhageni/***sv*Icterohaemorrhagiae (preferred hosts, peri-domiciliary rats), ***sv*Lai** (field mice), *sv*Hardjo-prajitno (cattle) and ***sv*Canicola/***sv*Pomona (dogs and pigs, respectively) were distinguishable via T_m_’s. As expected, T_m_’s were species-dependent (reflecting species-variable GC%), with *interrogans*/*kirschneri*/*noguchii* alleles/serovars producing T_m_’s ranging from 80 – 81°C; *borgpetersenii*, 81.5 – 82.5°C; *weilii*, 83°C; and *santarosai*, >84°C (Table 2 and **Appendix S1**), allowing joint inference of species and serotype, ergo *interrogans sv*Hardjo-prajitno could be distinguished from *borgpetersenii sv*Hardjobovis. Though not the recommended default, all nine sequence variants could be differentiated at 0.1°C-resolution (not shown). As we are conducting prospective field studies in both Peru and Sri Lanka, we also analyzed reference strains obtained from both regions. All strains originating from Sri Lanka (***sv*Geyaweera/***sv*Weersinghe, ***sv*Ratnapura, *sv*Ceylonica** and ***sv*Alice**) could be reliably differentiated from each other (80.6, 80.4, 81.6, 85°C), and from ***sv*Copenhageni/***sv*Icterohaemorrhagiae (80.2°C). By contrast, Peruvian reference strains originating from outside Iquitos, specifically those belonging to *L. santarosai*, amplified inconsistently, precluding T_m_ assignment and suggesting that further optimization might be necessary to reliably detect these serotypes.

### Serovar resolution using high-precision magnetic induction thermal cycling

Owing to differing instrument design philosophies, MIC-derived T_m_’s were on average ∼1°C higher (***σ*** ≤ 0.05) than those produced by the CFX96. Nonetheless, statistical comparison of the means showed that, as anticipated, the superior resolution of MIC improved allele resolution (up to 100 potential distinct T_m_’s at 0.05°C-resolution, and 200 at 0.025°C-resolution). Three previously unresolved (but distinct) alleles present in *sv*Bataviae, ***sv*Copenhageni/***sv*Icterohaemorrhagiae, and ***sv*Cynopteri**, could be assigned a T_m_ by MIC, **81.19±0.005°C, 81.22±0.000°C**, and **81.28±0.01°C**, respectively (Table 2). Alleles found in ***sv*Hardjo-prajitno/***sv*Wolffi and ***sv*Mankarso** could also be resolved by MIC, as were four distinct alleles present in ***sv*Australis/***sv*Bratislava*/sv*Djasiman/*sv*Grippotyphosa/*sv*Pyrogenes **81.87±0.004°C**; *sv*Autumnalis, **81.88±0.004°C**; ***sv*Canicola/***sv*Pomona, **81.83±0.010°C**; and *sv*Manilae, **81.79±0.010°C** (Table 2). Overall, all nine sequence variants present in *interrogans*/*kirschneri* serovars could be distinguished with strong statistical support. Some, such as *sv*Lai, ***sv*Copenhageni/***sv*Icterohaemorrhagiae and *sv*Mankarso, belonging to the same **Serogroup** or representing multi-species serovars, e.g., *sv*Hardjo subtypes prajitno and bovis.

### Reliability, sensitivity and specificity of *11108*-based assay vs clinical samples

During the five-year-period December 2002 – June 2007, 144 *Leptospira* strains were isolated from humans (50), peridomiciliary rats (34), livestock (32), and wild animals (28) in Iquitos and surrounding areas. The majority (65.3%) were either *interrogans, kirschneri* or *noguchii*. Fifty (34.7%) were identified as *santarosai*, with 27 (or 54%) derived from livestock (i.e., buffaloes, cattle and pigs) representing 84.4% of all isolates obtained from these sources. By contrast, only 14 human isolates were classified as *santarosai* (28%), as were six from wild animals (21.4%), two from rats (5.9%), and none from bats or dogs. To test the reliability and accuracy of our optimized 11108-based NAAT, we analyzed a sample of these isolates— inclusive of all species and **Serogroups** found in the region. Twenty-three were taxonomically classified as *interrogans*, 13 as *santarosai*, seven as *L. noguchii* and two, as *L. kirschneri*. Table 3 summarizes the 11108 qPCR-assay results of all human and rat isolates analyzed.

Based on CAAT (and/or PFGE-fingerprinting), 23 *interrogans* strains belonged to either serogroup **Icterohaemorrhagiae** (with all but two definitively typed as ***sv*Copenhageni**) or **Canicola** (***sv*Canicola**, and two closely related but seemingly distinct strains, BEL039 and CBC1203R). All produced consistent T_m_’s which were verified by BLAST. And so far, only two strains isolated in Iquitos, HAI1536 and MOR069, have been shown to have similar 11108 amplicons but different CAAT or PFGE fingerprints among the 19 analyzed that were classified as *interrogans* / *kirschneri* / *noguchii*. Similarly, isolate MMD4803—presumed distinct based on PFGE fingerprinting, could not be resolved from either of two HAI1378-derived isolates, nor CBC621. Overall, of 33 isolates analyzed, only two produced **11108**-inferred IDs that disagreed with CAAT or PFGE-based assignments, indicating correct inferences approximately ∼94% of the time, regardless of isolation date or source (human, clinical vs mammalian reservoir). Species inferences were infallible, and the assay correctly differentiated alleles/sequence variants 100% of the time. As is the case for most ‘exotic’ serovars, sequences were not publicly available for **Bataviae** strains, VAR132 (80.4°C), MOR069 (80.6°C), nor **Shermani** strains, HAI1378(U) and CBC621 (84.8°C), precluding definitive serovar assignment, but these were all confirmed via BLAST best-hits to be distinct and derived from 11108-amplicon variants grouping with other serotypes from the expected species, i.e., *noguchii* and *santarosai*, respectively. This, based upon T_m_-defined species-ranges. Taken together these findings indicate that the 11108-assay is highly reliable and capable of producing consistent results with strains isolated several years apart, originating from various hosts. And whereas T_m_ values vary based upon hardware/software options utilized (highlighted in Table 2), our approach consistently distinguished sequence variants.

Pilot experiments using archived culture-positive patient serum-samples from Sri Lanka (**12 isolates**, age range **=** three to five years-old) and Iquitos (**25**, range 11 – 15) produced mostly consistent results. Six of the recently archived samples amplified consistently across replicates and produced expected T_m_’s with respect to their cognate isolate. But six failed to amplify (Figure 1). Amplification of the older samples was less consistent, though all were positive with amplification evident in at least one replicate in all runs. And regrettably, most produced indefinite or quasi-stable T_m_’s (±0.1°C), though some, e.g., serum-sample PAD451 (81.0°C), produced reliable T_m_’s agreeing with CAAT-based classification of its cognate isolate, indicating that further refinement of our assay could allow retrospective analysis of bio banked serum-samples.

**Figure 1:**
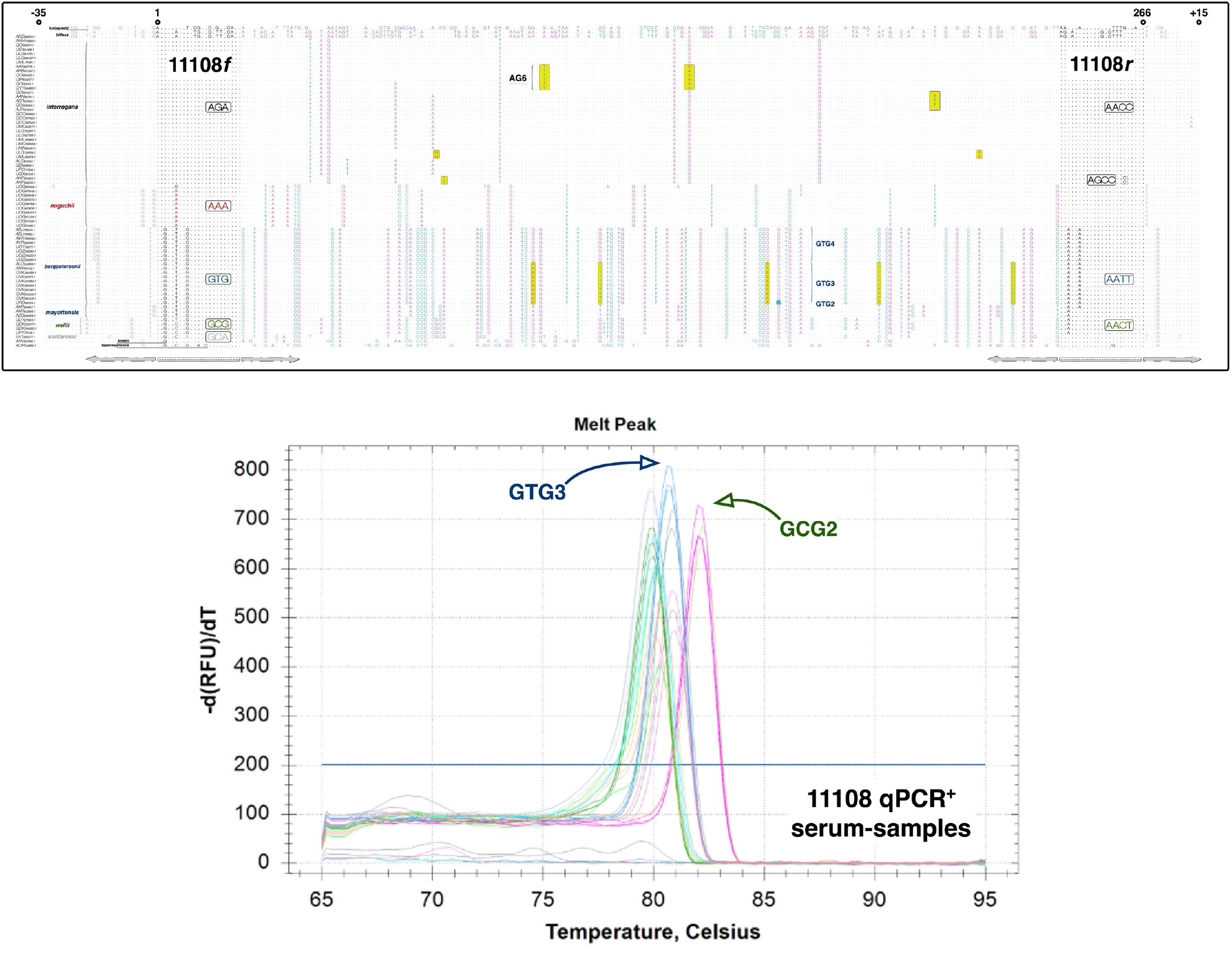
Schematic showing predicted and experimentally validated 11108-amplicon sequence-variants in patient sera. **Top Panel.** Global alignment of 266-bp region amplified by 11108 primer pairs **w/**additional nucleotides at the 5’ and 3’ end, (+35 and −15, respectively). Forward (i.e., 11108f) and reverse primers and their species-associated sequence variants, e.g., 11108f_**AGA** for *interrogans*/*kirschneri* and *noguchii* strains, have been indicated. *<u>Arrows</u>* below MSA indicate alternate primer sites. Examples of allele defining SNPs are enclosed in yellow boxes. **AG6** = *sv*Copenhageni. **GTG3** = *borgpetersenii*, unknown (**UNK**) **serogroup** (i.e., no agglutination in serotyping reactions). **GCG2** = *weilii* **UNK** serogroup. **AG6** is dominant human infecting strain in Iquitos (Peru), whereas **GTG3** and **GCG2** comprise novel non-*interrogans* serovars isolated from hospitalized patients in Sri Lanka—**Appendix S1**. *Leptospira* spp. shown: *kobayashi* (top, non-pathogen), *biflexa* (non-pathogen), *interrogans, noguchii, borgpetersenii, mayottensis, weilii, santarosai, kmetyi* and *tipperaryanensis* (bottom). **Bottom Panel**. CFX96-generated melt-curves (**w/o** Precision Melt Software), showing T_m_’s and allele assignments from six (of 12) qPCR-positive serum-samples. Three species were reliably distinguished: *interrogans* (unlabeled peak), *borgpetersenii* GTG3 and *weilii* GCG2, consistent with predicted allele assignments and 11108-based amplification of cognate isolates.

The sensitivity (true positives detected) was clearly superior to the *lipL32*-based assay used for comparison while maintaining 100% specificity.

## Discussion

Recognizing the public health importance of real-time serovar ID, we developed a novel single-tube NAAT capable of providing reliable *Leptospira* serovar predictions directly from diverse sample types. Rigorous validation against dog urine samples, proved our assay to be highly specific for group I pathogenic *Leptospira*, with no detectable off-target amplification of even genetically related (but usually hypo-virulent) group II species or of unrelated, poorly characterized microorganisms in complex microbial species-consortia. It dramatically outperformed a mainstream *lipL32*-based alternative and proved unique in its capacity to provide joint species and serotype IDs, while consistently detecting fewer than 10 genome equivalents/sample in serum and urine. From a public health perspective, the early successes of our approach provide pivotal proof-of-concept validation that should spur development of a newer class of NAATs specifically for the detection and direct identification of *Leptospira* serovars in varied clinical settings and epidemiological contexts.

The MAT was formerly the gold standard for leptospirosis diagnosis. But, owing to well-documented limitations (and antibody-based-diagnosis in general, it has been steadily supplanted by a bevy of NAATs, with *lipL32*-based real-time qPCR assays now routine. But, until now, NAATs by design could not generate epidemiologically useful information. And, while true that some 16S rRNA-based assays yield species IDs (18), these alone are meaningless from a public-health standpoint, as many important but distinct serovars belong to the same species, making it nigh impossible to assess the relative importance of potential reservoir species, domesticated or otherwise, in the maintenance and transmission of human leptospirosis. Furthermore, as diagnostic tests, 16S rRNA-based assays, though sensitive, are mostly genus-inclusive (and not specific for pathogenic *Leptospira*); they tend to produce false-positive results, and are practically unusable with certain sample types, such as urine (31). By contrast, our assay and *lipL32*-based alternatives are highly specific for pathogenic species, but unlike the latter, ours proved to be as sensitive as those targeting the 16S rRNA gene. And, of the three design philosophies, only our assay yielded joint predictions of infecting species and serovar. So, based on its validated sensitivity and specificity, empirically determined detection limit, and the verified accuracy of its predictions, this new NAAT holds considerable promise for real-time serovar ID, an important milestone for public health leptospirosis surveillance programs.

We also introduce the concept of serotype-specific T_m_’s (i.e., melt-temperatures) and show that under optimal conditions all unique 11108-amplicon variants could be reliably differentiated by this metric alone. More importantly, assay predictions proved infallible (i.e., unexpected T_m_’s were all well-supported by SNPs in sequenced amplicons), a remarkable improvement on the MAT, which is currently the only other diagnostic test permitting inference of infecting serovar. Sadly though, MAT-based predictions are mostly inaccurate (32, 33) or ambiguous (34), though presumably improved through analysis of paired (acute and convalescent) sera. As far as we know, the two most medically important serovars worldwide, *L. interrogans sv*Copenhageni and *sv*Canicola, were easily and reproducibly distinguished, as were others with shared host preferences. Even multi-species serovars, such as *sv*Hardjo *interrogans* subtype prajitno and *borgpetersenii* subtype bovis, were easily distinguished. And novel or poorly characterized serovars with little to no genome information could be differentiated from each other and common serovars based on T_m_ and if needed, amplicon-derived sequence data, as was the case with buffalo isolate (CBC621). In addition, because of the region we’ve selected further improvements or customizations are not only possible but easily developed. Further, this is but one of many gene targets we’ve identified so far that appear to have clinical utility for rapid diagnoses and prognostic testing.

Our approach is not intended to replace specialized serotyping methods, such as MLST profiling or PFGE fingerprinting, which by design have superior resolving power, but rather to address a clear need for NAATs capable of rapid and reliable serovar ID directly from varied sample types. NAATs that are readily adapted to diverse clinical settings and public health surveillance initiatives. Additionally, in recent years emphasis has been placed upon bridging the gap between historical knowledge of *Leptospira* serological and molecular techniques, with growing concern that crucial epidemiological information will be lost in the transition to modern rapid diagnostics (35).

## Acknowledgements

Our thanks go out to the Lars Eckmann laboratory for their kind gift of negative control bacterial species, to the anonymous patients who contributed to our isolate bank, Lee Smythe and others at the Queensland Health Leptospirosis Reference Laboratory, and all others who contributed to this study.

## Approvals and Funding

This study was funded by the National Institute for Allergy and Immunology **U19AI115658, R21AI108276** and **R21AI164106**.

